# Random forest based similarity learning for single cell RNA sequencing data

**DOI:** 10.1101/258699

**Authors:** Maziyar Baran Pouyan, Dennis Kostka

## Abstract

**Motivation:** Genome-wide transcriptome sequencing applied to single cells (scRNA-seq) is rapidly becoming an assay of choice across many fields of biological and biomedical research. Scientific objectives often revolve around discovery or characterization of types or sub-types of cells, and therefore obtaining accurate cell–cell similarities from scRNA-seq data is critical step in many studies. While rapid advances are being made in the development of tools for scRNA-seq data analysis, few approaches exist that explicitly address this task. Furthermore, abundance and type of noise present in scRNA-seq datasets suggest that application of generic methods, or of methods developed for bulk RNA-seq data, is likely suboptimal.

**Results:** Here we present RAFSIL, a random forest based approach to learn cell–cell similarities from scRNA-seq data. RAFSIL implements a two-step procedure, where feature construction geared towards scRNA-seq data is followed by similarity learning. It is designed to be adaptable and expandable, and RAFSIL similarities can be used for typical exploratory data analysis tasks like dimension reduction, visualization, and clustering. We show that our approach compares favorably with current methods across a diverse collection of datasets, and that it can be used to detect and highlight unwanted technical variation in scRNA-seq datasets in situations where other methods fail. Overall, RAFSIL implements a flexible approach yielding a useful tool that improves the analysis of scRNA-seq data.

**Availability and Implementation:** The RAFSIL R package is available online at www.kostkalab.net/software.html

## 1 Introduction

Sequencing transcriptomes of single cells (scRNA-seq) is becoming increasingly common, as technology evolves and costs decline. Studying gene expression genome-wide at single cell resolution overcomes intrinsic limitations of bulk RNA sequencing, where expression levels are averaged over thousands or millions of cells. scRNA-seq enables researchers to more rigorously address questions about the cellular composition of tissues, the transcriptional heterogeneity and structure of “cell types”, and how this may change, for instance during development or in disease (Kumar *et al.*, 2017; Patel *et al.*, 2014). Identifying group structure is therefore a crucially important step in most scRNA-seq data analyses, and it has yielded exciting discoveries of novel cell types and revealed previously un-appreciated sub-populations and heterogeneity of known types of cells (Kumar *et al.*, 2017).

Identifying group structure in scRNA-seq data is, however, not without challenges. Even for bulk RNA sequencing no gold standard has emerged in the field (Conesa *et al.*, 2016), and for single cell RNA sequencing several factors further complicate the task. These include additional biological heterogeneity induced by the inherent stochasticity of gene expression in single cells, and technical noise rooted in cell processing, cell lysis, and library preparation from extremely low amounts of “input” messenger RNA (Adam *et al.*, 2017). The latter, for example, leads to dropout events, where no RNA is measured for a gene actually expressed in a cell. It is estimated that 50–95% of a cell’s mRNA are not measured by current technologies (Adam *et al.*, 2017; Svensson *et al.*, 2017). While the relative magnitude of such factors will depend on the specific technology used, it is fair to assume they play a role in most, if not all, scRNA-seq studies. Therefore, there is a need for computational approaches that take the specific nature of scRNA-seq data into account and enable researchers to accurately and reliably identify, visualize, and explore group (or population) structure of single cells. To address that need we developed RAFSIL, a random forest based method for learning similarities between cells from single cell RNA sequencing experiments.

Related work includes clustering methods, which implicitly or explicitly rely on a similarity concept and are commonly used to group objects. Examples of approaches developed specifically for scRNA-seq data include the combination of Pearson correlation with robust k-means clustering (Grün *et al.*, 2015), and the use of consensus clustering (Strehl and Ghosh, 2002) to obtains stable cell groupings by Kiselev *et al.* (2017b). Žurauskienė and Yau (2016) combine agglomerative clustering with principal component analysis, while Lin *et al.* (2017) explore the use of neural networks (Hagan *et al.*, 1996) for clustering and dimension reduction. More closely related to our work is SIMLR (Wang *et al.*,2017b), an approach based on multiple kernel learning (Lanckriet *et al.*,2004) that directly learns similarities between single cells. However, SIMLR is built around a clustering paradigm, and the user is asked to provide the algorithm with a specific cluster number to guide similarity learning.

In contrast, RAFSIL similarities are based on random forests (Breiman, 2001), and our approach requires no prior information about group structure. We show RAFSIL learns similarities that faithfully represent group structure in scRNA-seq data; when used for dimension reduction and clustering they provide an accurate visualization of datasets and enable exploratory analyses for cell type identification and discovery. Importantly, RAFSIL compares favorably with the current state-of-the-art showing high accuracy and robustness, and we demonstrate how it enables the identification of technical variation that remains hidden with other approaches.

## 2 Methods

We assume normalized gene expression data on log-scale of *n* cells for *p* genes is available, organized into a p × n expression matrix **X** = (***x***_1_, ***x_2_,…, x***_n_), where ***x***_i_ indicates the expression of *p* genes in cell *i* ***x_i_*** = *(x_i1_,x_i2_,…,x_ip_)*^ʹ^.

### 2.1 Gene filtering

We consider three types of gene filters for the scRNA-seq data matrix **X**:

> *All genes (ALL).* All genes in ***X*** are considered that have non-zero expression in at least one cell in the dataset. This is the most inclusive set of genes.
>
> *Frequency filtering (FRQ).* Here we consider only genes that are expressed in a certain fraction of cells. Specifically, we choose 6%, as reported by (Kiselev *et al.*, 2017b) for our analyses.
>
> *Highly expressed genes (HiE).* The subset of frequency-filtered genes is further narrowed down to consider genes with “high” expression across cells. In each cell, expressed genes are sorted in decreasing order of expression and the top 10% are marked as highly expressed. To focus on genes that are frequently highly expressed across cells, we discard half of the genes that are highly expressed in the fewest cells This approach yields a set of genes that are highly expressed across cells, but still allows for variability in gene expression.

In the following, we describe our approach for <u>ra</u>ndom <u>f</u>orest based <u>si</u>milarity <u>l</u>earning (RAFSIL) from scRNA-seq data. We developed two methods, RAFSIL1 and RAFSIL2, which are both two-step procedures. They share a feature-construction step and then apply different types of random forest (RF) based similarity learning.

### 2.2 RAFSIL: Feature construction

***RAFSIL gene filtering and clustering.*** For the RAFSIL methods, we apply the frequency filter described above, and then derive gene clusters as follows: First, principal component analysis is applied to the gene-filtered expression matrix ***X*** (treating genes as observations and cells as features), and we keep the most informative principal components as selected by the “elbow method” Thorndike (1953). Next, we apply k-means clustering (`kmeans++`, Arthur and Vassilvitskii (2007); Mouselimis (2017)) to this reduced representation of genes and derive gene clusters, where we determine the number of clusters by finding the elbow point of the sum of squared errors as a function of increasing cluster numbers. This yields a partition of frequency-selected genes into *k* disjoint clusters.

***RAFSIL Spearman feature space construction.*** Gene clustering decomposes the column space of ***X*** into orthogonal sub-spaces, and we characterize all cell based on its similarities with all other cells in each sub-space. Specifically, and we calculate *n × n* cell-cell similarity matrices {*C*_1_,…,*C_k_*} using Spearman rank correlation and genes restricted to the respective clusters derived beforehand. For each similarity matrix *C_i_* we perform PCA, and again keep *m_i_* informative principal components identified by the elbow method. This yields *k* matrices 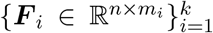 based on genes in cluster *i*, where each cell is embedded by its principal components derived from local similarities (i.e., similarities calculated using only genes in a gene cluster). We then construct a final feature matrix ***F*** by juxtaposing matrices from individual gene clusters:

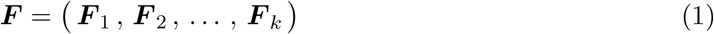

The number of columns in ***F*** (i.e., the number of features 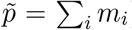) is data-dependent, and each cell i is now described by a feature vector 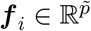 (the *i*-th row of ***F***). In the following we use these features for random forest based similarity learning.

### 2.3 RAFSIL: RAndom Forest based Similarity Learning

Random Forests (RFs) are an established classification method based on ensembles of decision trees (Breiman, 2001). However, they can also be used in an unsupervised setting to infer similarities between objects (Breiman and Cutler, 2003

#### 2.3.1 RAFSIL1

Here we describe an approach for RF based similarity learning (Breiman and Cutler, 2003; Shi and Horvath, 2006) that has been applied to various types of biomedical data (Seligson *et al.*, 2005; Ramirez *et al.*, 2017) and is implemented in the `randomForest` package for the **R** programming language (Liaw and Wiener, 2017). In (Pouyan and Nourani, 2017) the RAFSIL1 approach (without the feature construction step) was applied to Cytometry by Time of Flight (CyTOF) data, where protein expression of several marker genes (typically less than 50) is assessed.

Next, we briefly summarize RF based similarity learning: To cast the unsupervised similarity learning problem into a problem suitable for RFs, a “synthetic” dataset is generated, for instance by randomly shuffling the values of each feature independently; then, an RF classifier is trained to distinguish the shuffled data from the un-shuffled data (***F*** in our notation). Let ***fi*** denote the *i*-th row of ***F***. If we assume the RF classifier contains *N* trees and define `nt`(***fi***, ***fj***) as the number of trees that classify cells ***fi*** and ***fj*** via the same leaf, then the RF based *n* **×** *n* similarity matrix **S** is defined via *Sij* = nt(***fi***, ***fj***)/N. A corresponding dissimilarity matrix D can then be obtained via D_ij_ = 1 – *S_ij_*. In the following we use the term similarity and dissimilarity interchangeably, referring to ***S*** and ***D***, respectively. Repeating this procedure *B* times allows us to aggregate individual similarity matrices ***S^i^*** into a final matrix 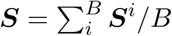 and corresponding ***D***. We used *B* = 50 for our experiments.

#### 2.3.2 RAFSIL2

We now describe how we use the RF classifier to construct (dis)similarity matrices without the need for synthetically generated datasets. The general idea, as in the above method, is to exploit feature dependence. However, we proceed as follows: After selecting a single feature *j* (the *j*–th column of the feature matrix **F**) we quantize its values to derive class labels 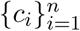 for all cells. We use partitioning around medoids as implemented by the `pamk` function provided by the R package `fpc` (Hennig, 2018), which also estimates the optimal number of clusters. Then, remove the *j*-th column from **F** and use the RF classifier to learn the obtained class labels with this reduced dataset. The resulting random forest than yields a similarity between cells as described above. Repeating this procedure for all features yields 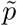 random forest classifiers with corresponding similarity measures ***S****^i^*, and averaging as described for RAFSIL1 above results in a final pair of similarity and dissimilarity matrices ***S*** and ***D***, respectively. As before, we use the `randomForest` package for R (Liaw and Wiener, 2017) with its default forest size of 500 trees.

### 2.4 Performance evaluation

To evaluate our approach, we apply RAFSIL1/2 to ten scRNA-seq datasets that have pre-annotated cell populations, and we compare results with current state of the art approaches. We distinguish three different scenarios, namely similarity learning, dimension reduction and clustering. All of these play critical roles in exploring, visualizing and interpreting scRNA-seq data, but they have different objectives and we evaluate them accordingly.

#### 2.4.1 Similarity learning

For similarity learning, we compare our method with SIMLR (Wang *et al.*, 2017b), the only scRNA-seq method that advertises similarity learning. In addition, we explored common similarity/dissimilarity measures: Euclidean distance, Pearson and Spearman correlation, applied to the full (ALL), frequency-filtered (FRQ) and highly-expressed (HiE) sets of genes (see Section 2.1 for details on the gene sets). Following Wang *et al.* (2017b) the metric we choose to evaluate similarity learning is the nearest neighbor error (NNE) Van Der Maaten *et al.* (2009). The NNE is calculated by using a nearest neighbor classifier based on the target similarity to be evaluated: For a given set of labeled cells, an unlabeled cell is classified with the same label as its most similar labeled neighbor. Predictions for each cell are obtained via 10-fold cross-validation (CV), and the NNE then reports the fraction of mis-classified cells. Because in the 10-fold CV procedure data are randomly split into 10 folds (9 for training, 1 for validation) we report averages over 20 runs. The NNE is a direct reflection of how well the learned dissimilarity measure captures the pre-annotated class labels. For SIMLR we used the SIMLR R package (Wang *et al.*, 2017a), provided it with all genes (ALL) and evaluated the similarity matrix returned by the `SIMLR` function with default options. For SIMLR we needed to provide the option `normalize=TRUE` for the Treutlein dataset, otherwise the program would abort. We have indicated this by putting the respective values in parentheses in the relevant result tables.

#### 2.4.2 Dimension reduction

To evaluate the results of dimension reduction, we use the same NNE metric as for evaluating similarity (see above), but in this case applied to the reduced dimensional projection. That is, we first perform similarity learning. Then we use the resulting similarity matrix as input for a dimension reduction algorithm, which sees each cell as a vector of its’ similarities. Finally we calculate the NNE based on Euclidean distance in the reduced-dimensional space.

For all methods we choose two as the number of dimensions to project down to, and we compared the following approaches for dimensionality reduction: t stochastic neighbor embedding (tSNE, van der Maaten and Hinton (2008)), principal component analysis (PCA), and probabilistic PCA (pPCA, Tipping and Bishop (1999)). We also skip the similarity learning step and directly apply dimension reduction to cells characterized by their highly expressed genes (Data-HiE-in Table 3). For probabilistic PCA we used the implementation provided by the `pcaMethods` R package (Stacklies *et al.*, 2007; Kiselev *et al.*, 2017a) and for t stochastic neighbor embedding the `Rtsne` R package (Krijthe, 2015). We used tSNE with default values for all datasets except Treutlein, where we set the perplexity to 20.

#### 2.4.3 Clustering

We also evaluate the performance of RAFSIL1/2 in the context of clustering; that is, we ask how well group structure inferred based on RAFSIL1/2 similarities agrees with pre-annotated cell populations. This allows us to expand the methods we compare RAFSIL with, because in addition to the approaches we compared with for similarity learning and dimension reduction, we can now add algorithms that have no explicit similarity learning step. Specifically, we add SC3 (Kiselev *et al.*, 2017b,c), pcaReduce (Žurauskienė and Yau, 2016, 2015) and SINCERA (Guo *et al.*, 2015; Guo, 2017) to our comparisons. These methods, and SIMLR, are geared towards scRNA-seq clustering, and we provide each method with the number of pre-annotated populations for each dataset and the expression profiles comprising the complete set of expressed genes (ALL).

***Clustering methods.*** For RAFSIL1/2 and Spearman correlation we implemented two clustering strategies. First, using similarities as a vector embedding for each cell, we run k-means clustering (KM) to infer group labels. Second, we perform hierarchical clustering with average linkage (HC) using learned dissimilarities (1 – *ρ* for Spearman correlation). For k-means clustering we use `kmeans++` as provided by the R package pracma (Borchers, 2017), while for hierarchical clustering we use the base functionality provided within R through the `stats` package (R Core Team, 2017). Like for the other methods, we set the number of clusters to the known number of different cell labels (Kiselev *et al.*, 2017b,c).

***Evaluation metric.*** To evaluate clustering results we calculate two performance metrics: the adjusted Rand index (ARI) and normalized mutual information (NMI). Both of them are popular metrics to evaluate clustering results in the context of a known labeling in single cell data (Wang *et al.* (2017b); Kiselev *et al.* (2017b); Hubert and Arabie (1985); Vinh *et al.* (2010)). The ARI is defined as follows: Assume we cluster *n* cells into *k* clusters. Let 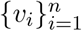 denote the inferred cluster labels, and 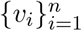 the pre-annotated labeling. Then

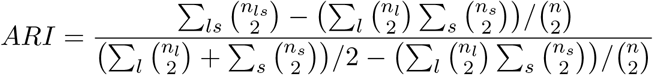

where *l* and *s* enumerate the *k* clusters, and 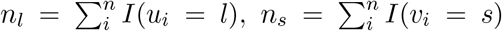 and 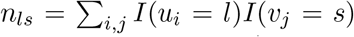 with I(x = y) the indicator function that is one for *x* = *y* and zero otherwise. The ARI is one if the inferred labels correspond perfectly to the known labels, and it decreases with increasing disagreement.

For the normalized mutual information, let p*_i_* = n*_i_*/n and q*_s_* = n*_s_*/n ans z*_ls_* = n*_ls_*/n. Then h(*u*)=Σ*_l_p_l_* log(*p_l_*) and h(*v*) = Σ*_s_ q_s_* log(*q_s_*) are the respective entropies of the two clusterings, and *i*(*u, v*) = Σ_l, s_ z_is_ log(z*_ls_*/p*_l_*/q*_s_*) is their mutual information. The normalized mutual information is then defined as 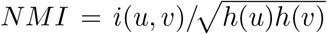. Like the ARI the NMI is one for perfectly overlapping clusterings, and it decreases with increasing disagreement. It is bounded by zero from below. For ARI and NMI we report median values over 20 clustering runs in our clustering evaluation.

***Clustering in reduced dimensions.*** We also evaluate clustering results after dimension reduction. To do so, we build on the results from evaluating dimension reduction with the nearest neighbor error (see Section 2.4.2). For each similarity learning approach we assess the corresponding dimension reduction method with the smallest NNE and then perform standard k-means and hierarchical clustering in reduced dimensions. Results are then evaluated as described above. However, here we use Pearson correlation and not Spearman correlation as a representative for generic similarity learning, because it performs slightly better (see Table 3).

### 2.5 Data used and software availability

Datasets used in the majority of our analyses are summarized in Table 1. Patel, Pollen, Goolam and Treutlein datasets were downloaded from https://hemberg-lab.github.io/scRNA.seq.datasets/; Usoskin, Buettner and Kolod datasets were downloaded from https://github.com/BatzoglouLabSU/SIMLR. The Engel and Lin datasets can be found in the supporting material of Lin *et al.* (2017) and were downloaded from http://128.2.210.230:8080/; the label “Lin” in our result tables refers to the combination of three primary datasets described in the Methods section there. Finally, the Leng dataset was obtained from https://bioinfo.uth.edu/scrnaseqdb/.

**Table 1:** List of datasets analyzed and their attributes

| Dataset | # cells | # genes | # populations | sparsity (in %) | Units | Reference |
| --- | --- | --- | --- | --- | --- | --- |
| Patel | 430 | 5,948 | 5 | 0 | TPM | (Patel *et al.*, 2014) |
| Buettner | 182 | 9,573 | 3 | 37 | FPKM | (Buettner *et al.*, 2015) |
| Engel | 203 | 21,690 | 4 | 80 | TPM | (Engel *et al.*, 2016) |
| Kolod | 704 | 13,473 | 3 | 10 | CPM | (Kolodziejczyk *et al.*, 2015) |
| Goolam | 124 | 41,480 | 5 | 69 | CPM | (Goolam *et al.*, 2016) |
| Usoskin | 622 | 17,772 | 4 | 78 | RPM | (Usoskin *et al.*, 2015) |
| Treutlein | 80 | 23,271 | 5 | 90 | FPKM | (Treutlein *et al.*, 2014) |
| Leng | 460 | 19,084 | 4 | 47 | TPM | (Leng and Kendzierski, 2015) |
| Pollen | 301 | 9,966 | 11 | 67 | TPM | (Pollen *et al.*, 2014) |
| Lin | 402 | 9,437 | 16 | 43 | TPM | (Lin *et al.*, 2017) |

For our analysis underlying Figure 1, the Usoskin and Kolod datasets were re-downloaded to obtain normalized expression values without batch corrections. For Usoskin, we downloaded this information from the “External resource Table 1”, available at http://linnarssonlab.org/drg/; for Kolod, data was downloaded from https://www.ebi.ac.uk/teichmann-srv/espresso/.

**Figure 1:**
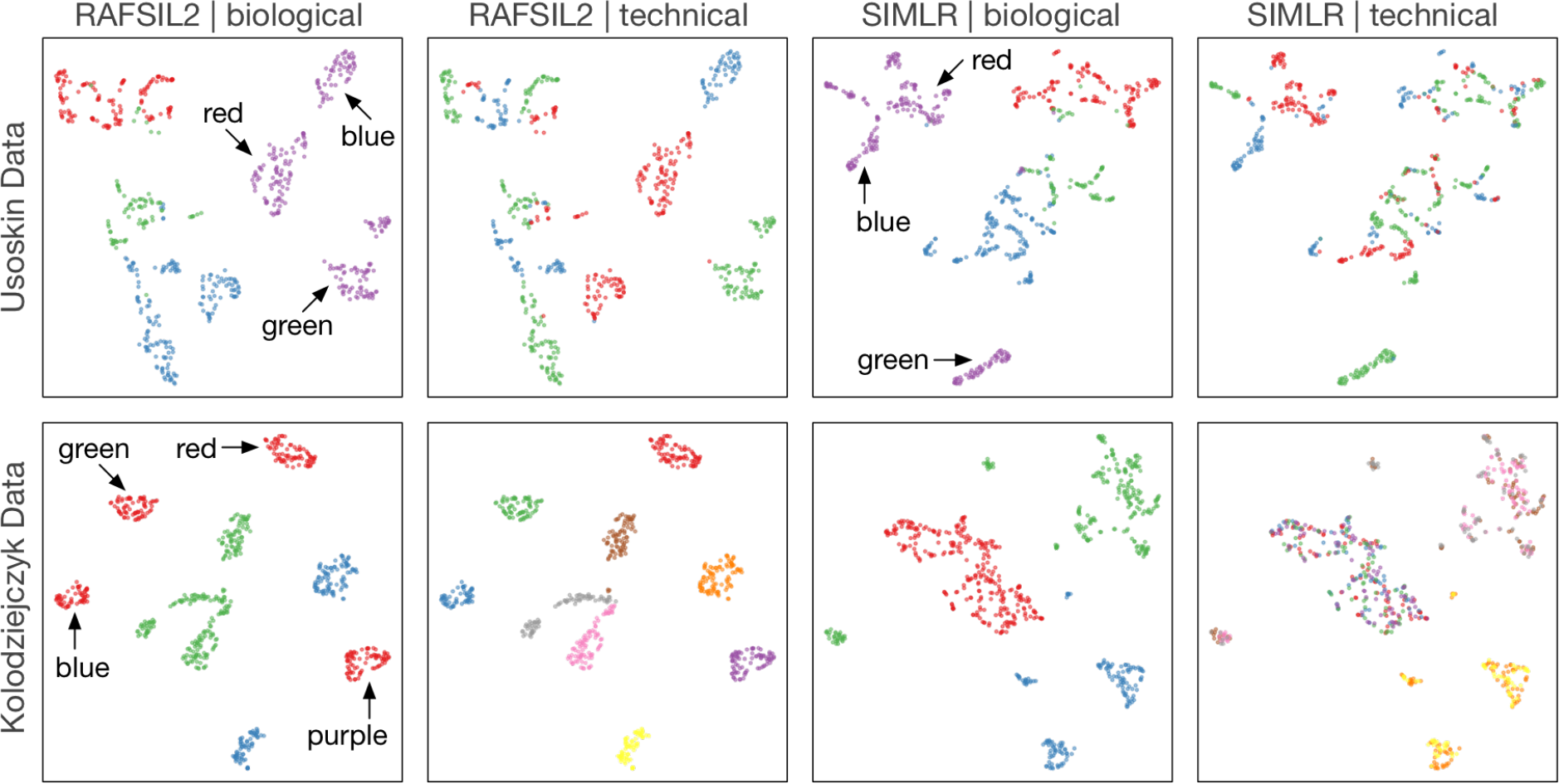
RAFSIL2 discovers unwanted variation. This figure shows tSNE plots for two datasets: Data from Usoskin *et al.* (2015) in the first row, and from Kolodziejczyk *et al.* (2015) in the second row. Cells are colored according to biologically meaningful annotations in panels one and three, and according to technical covariates in panels two and four. In the first row we see that sub-structure in biologically meaningful groupings can be explained through technical variables for both methods. In the second row this still holds true for RAFSIL2, but SIMLR does not highlight the unwanted technical variation present in the data. For more details see Section 3.2.2.

The `RAFSIL R` package is available at www.kostkalab.net/software.html.

## Results

### 3.1 A random forest based approach for single cell similarity learning

Here we present RAFSIL, a random forest (RF) based approach for learning similarities from single cell RNA-sequencing data. Random forest based similarity learning (Shi and Horvath, 2006) is a way to apply random forests (Breiman, 2001) to unsupervised learning and derive similarities between objects (Shi and Horvath, 2006; Breiman and Cutler, 2003). In particular, RF based similarity learning is robust to outliers and has built-in feature selection, which is appealing for analyzing high-dimensional and noisy data, like single cell RNA sequencing profiles. We also note that this approach is fundamentally different from ensemble approaches working with multiple clusterings of a dataset, see (Yan *et al.*, 2013, Section 3). To apply RF based similarity learning to single cell RNA sequencing (scRNA-seq) data, we implemented an approach we call RAFSIL. It is a two-step procedure, where in the first step we pre-process scRNA-seq expression data (feature construction step) and in a second step then perform RF-based similarity learning (similarity learning step).

The feature construction step is a heuristic approach designed to deal with the noise and sparsity typically present in scRNA-seq data (Yuan *et al.*, 2017). Briefly, we first find an orthogonal sub-space decomposition of the input space of cells, and then we describe each cell by its “local” similarities to other cells in each sub-space separately, which we then aggregate to a final feature set. Details on the feature construction step are in Section 2.2.

For the RF-based similarity learning step we explore two different approaches: RAFSIL1 and RAFSIL2. RAFSIL1 is a straight forward application of the methodology of Shi and Horvath (2006) to learn similarities between single cells described by the features recovered in our feature construction step. The general idea is to use random forest to discriminate between the real and a synthetic dataset, where the latter is derived from the real data by applying perturbations that destroy feature correlations. Similarity between cells is then quantified by co-classification of pairs of cells via the same leaf across trees in the random forest. For RAFSIL2, we apply random forests to unsupervised learning in a different way. For each feature, we quantize its values to derive class labels for cells, and then use the other features to predict these labels using a random forest. Similarity is then quantified in the same way as described before. Details about RAFSIL1 and RAFSIL2 are in Sections 2.3.1 and 2.3.2.

In the following we show that RAFSIL1/2 compare favorably with current approaches across a variety of scenarios. We also show how the method enables identification of unwanted technical variation in scRNA-seq datasets.

### 3.2 Similarities learned by random forests accurately characterize single cell RNA sequencing data

We applied RAFSIL1 and RAFSIL2 to a diverse collection of single cell RNA sequencing datasets (Table 1) and compared their performance with state-of-the-art approaches. In our analyses we distinguish three scenarios: Similarity learning, dimension reduction, and clustering. For similarity learning, we evaluate how well inferred pairwise similarities characterize pre-annotated cell populations (i.e., class labels fore cells). For dimension reduction, we use the inferred similarities as features and project each cell into two dimensions. We then evaluate how well the resulting euclidean distances between projected cells characterize pre-annotated cell populations. Finally, we also evaluate how accurately inferred similarities allow clustering algorithms to reproduce available class labels; we apply clustering algorithms to two settings: the originally inferred similarities, and similarities in reduced-dimensional projections inferred by dimension reduction approaches.

#### 3.2.1 Similarity learning

***Random forest based similarities accurately capture cell population structure in scRNA-seq data.*** We applied our RAFSIL algorithms to ten datasets (see Table 1), where labels for cell populations have been pre-annotated. We assess the learned similarities in terms of the nearest neighbor error (NNE), which is the mis-classification rate of a nearest neighbor classifier (see Section 2.4.1 for details). We compare RAFSIL1/2 to SIMLR (Wang *et al.*, 2017b), which performs similarity learning specifically for scRNA-seq data, and to (dis)similarities as assessed by Euclidean distance, Spearman and Pearson correlation. For the latter three we assess three gene selection strategies: all expressed genes (ALL), frequently expressed genes (FRQ), and only highly expressed genes (HiE); see Section 2.4 for a more detailed description.

Results are summarized in Table 2. We see that RAFSIL1/2 and SIMLR learn similarities that accurately characterize annotated cell populations (i.e., they have low NNE). We also find that RAFSIL and SIMLR substantially outperform Euclidean distance and the two correlation-based similarities, and that RAFSIL2 shows the best overall performance. For the Euclidean distance and the correlation-based approaches we also observe that focusing on highly expressed genes improves performance for all of them.

**Table 2:**
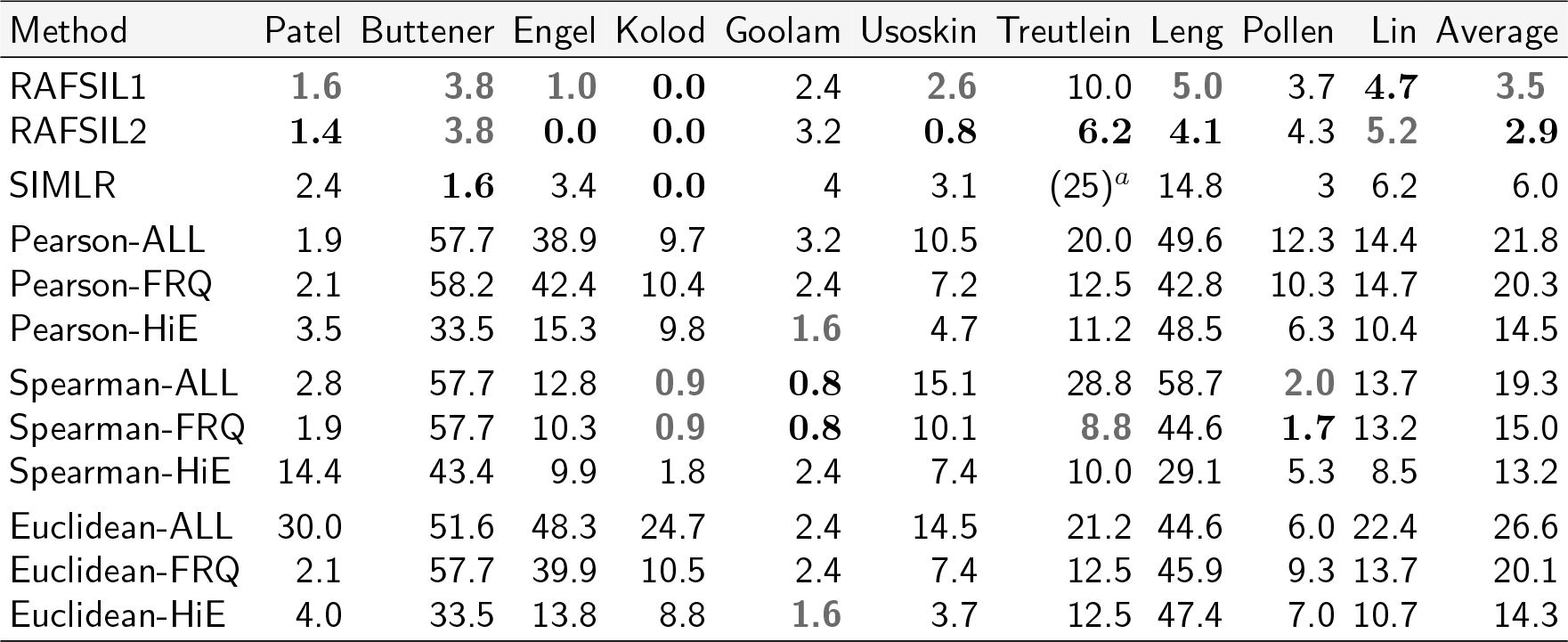
Nearest neighbor error values for similarity learning (in percent, lower is better). ALL = all expressed genes, FRQ = frequency-filtered genes, HiE = highly-expressed genes. *^α^* Parentheses indicate that SIMLR was run with different parameters for this dataset.

| Method | Patel | Buttner | Engel | Kolod | Goolam | Usoskin | Treutlein | Leng | Pollen | Lin | Average |
| --- | --- | --- | --- | --- | --- | --- | --- | --- | --- | --- | --- |
| RAFSIL1 | **1.6** | **3.8** | **1.0** | **0.0** | 2.4 | **2.6** | 10.0 | **5.0** | 3.7 | **4.7** | **3.5** |
| RAFSIL2 | **1.4** | **3.8** | **0.0** | **0.0** | 3.2 | **0.8** | **6.2** | **4.1** | 4.3 | **5.2** | **2.9** |
| SIMLR | 2.4 | **1.6** | 3.4 | **0.0** | 4 | 3.1 | (25)<sup>a</sup> | 14.8 | 3 | 6.2 | 6.0 |
| Pearson-ALL | 1.9 | 57.7 | 38.9 | 9.7 | 3.2 | 10.5 | 20.0 | 49.6 | 12.3 | 14.4 | 21.8 |
| Pearson-FRQ | 2.1 | 58.2 | 42.4 | 10.4 | 2.4 | 7.2 | 12.5 | 42.8 | 10.3 | 14.7 | 20.3 |
| Pearson-HiE | 3.5 | 33.5 | 15.3 | 9.8 | **1.6** | 4.7 | 11.2 | 48.5 | 6.3 | 10.4 | 14.5 |
| Spearman-ALL | 2.8 | 57.7 | 12.8 | **0.9** | **0.8** | 15.1 | 28.8 | 58.7 | **2.0** | 13.7 | 19.3 |
| Spearman-FRQ | 1.9 | 57.7 | 10.3 | **0.9** | **0.8** | 10.1 | **8.8** | 44.6 | **1.7** | 13.2 | 15.0 |
| Spearman-HiE | 14.4 | 43.4 | 9.9 | 1.8 | 2.4 | 7.4 | 10.0 | 29.1 | 5.3 | 8.5 | 13.2 |
| Euclidean-ALL | 30.0 | 51.6 | 48.3 | 24.7 | 2.4 | 14.5 | 21.2 | 44.6 | 6.0 | 22.4 | 26.6 |
| Euclidean-FRQ | 2.1 | 57.7 | 39.9 | 10.5 | 2.4 | 7.4 | 12.5 | 45.9 | 9.3 | 13.7 | 20.1 |
| Euclidean-HiE | 4.0 | 33.5 | 13.8 | 8.8 | **1.6** | 3.7 | 12.5 | 47.4 | 7.0 | 10.7 | 14.3 |

#### 3.2.2 Dimension reduction

***Dimension reduction improves similarity learning.*** We performed dimension reduction on the learned similarities obtained from RAFSIL1/2, and compared results with the same methods used in the previous section: SIMLR and Euclidean distance, as well as Spearman and Pearson correlation. We again use the NNE as a quality metric (on Euclidean distances in the reduced-dimensional space, for all methods), and results are summarized in Table 3. As a baseline approach we also included dimension reduction directly on the expression data (Data– in Table 3); this is different from the other methods, where we apply dimension reduction to cells described by their similarities with other cells (see Section 2.4.2).

**Table 3:** Nearest neighbor error values for dimension reduction (in percent, lower is better). tSNE = t stochastic neighbor embedding, PCA = principal component analysis, pPCA = probabilistic PCA. *^α^* Parentheses indicate that SIMLR was run with different parameters for this dataset.

| Method | Patel | Buttner | Engel | Kolod | Goolam | Usoskin | Treutlein | Leng | Pollen | Lin | Average |
| --- | --- | --- | --- | --- | --- | --- | --- | --- | --- | --- | --- |
| RAFSIL1-tSNE | **1.9** | 3.8 | **0.5** | **0.0** | 4.0 | **1.0** | 7.5 | **4.1** | **2.7** | **5.5** | **3.1** |
| RAFSIL1-PCA | 8.1 | 4.4 | 11.3 | **0.0** | 9.7 | 21.5 | 12.5 | 26.5 | 12.6 | 24.9 | 13.2 |
| RAFSIL1-pPCA | 7.7 | 4.4 | 11.3 | **0.0** | 9.7 | 22.5 | 15.0 | 25.4 | 12.3 | 24.4 | 13.3 |
| RAFSIL2-tSNE | **1.9** | **2.7** | **0.0** | **0.0** | 4.8 | **0.6** | **6.2** | **4.6** | 4.0 | **9.2** | **3.4** |
| RAFSIL2-PCA | 10.2 | 6.6 | 5.9 | **0.0** | 4.8 | 5.6 | 12.5 | 25.9 | 16.3 | 33.1 | 12.1 |
| RAFSIL2-pPCA | 9.8 | 7.1 | 4.9 | **0.0** | 4.0 | 5.3 | 11.2 | 26.3 | 14.3 | 30.6 | 11.4 |
| SIMLR-tSNE | 3.7 | 3.3 | 4.4 | **0.0** | 4.8 | 5.5 | (26.2)<sup>a</sup> | 19.8 | 3.0 | 15.7 | 8.6 |
| SIMLR-PCA | 6.7 | **2.2** | 27.1 | **0.1** | 11.3 | 6.4 | (43.8) | 36.3 | 22.9 | 51.0 | 20.8 |
| SIMLR-pPCA | 7.4 | **2.2** | 27.6 | **0.1** | 9.7 | 5.9 | (45) | 37.0 | 22.3 | 53.2 | 21.0 |
| Data-HiE-tSNE | 7.4 | 12.1 | 14.3 | 0.3 | 1.6 | 3.7 | 15.0 | 37.0 | 3.3 | 10.7 | 10.5 |
| Data-HiE-PCA | 40.7 | 25.8 | 13.3 | 1.4 | 4.8 | 34.1 | 31.2 | 56.1 | 16.3 | 40.5 | 26.4 |
| Data-HiE-pPCA | 40.5 | 28.6 | 14.3 | 1.4 | 7.3 | 33.3 | 32.5 | 57.4 | 17.3 | 41.5 | 27.4 |
| Euclidean-HiE-tSNE | 4.4 | 4.4 | 3.9 | 0.4 | 6.5 | 8.2 | 23.8 | 39.1 | 5.3 | 21.1 | 11.7 |
| Euclidean-HiE-PCA | 36.5 | 7.7 | 35.0 | 7.0 | 25.8 | 58.7 | 32.5 | 52.8 | 19.9 | 39.1 | 31.5 |
| Euclidean-HiE-pPCA | 36.0 | 8.8 | 39.4 | 6.8 | 28.2 | 57.7 | 32.5 | 53.0 | 20.3 | 38.8 | 32.2 |
| Pearson-HiE-tSNE | 2.8 | 9.3 | 3.0 | **0.0** | **1.6** | 2.1 | 17.5 | 24.1 | **2.3** | 20.1 | 8.3 |
| Pearson-HiE-PCA | 25.1 | 23.1 | 16.3 | **0.1** | 2.4 | 27.8 | 12.5 | 49.1 | 10.6 | 27.9 | 19.5 |
| Pearson-HiE-pPCA | 24.0 | 23.1 | 17.2 | 0.3 | 2.4 | 27.5 | 15.0 | 47.6 | 11.3 | 28.1 | 19.7 |
| Spearman-HiE-tSNE | 3.3 | 11.0 | 1.0 | **0.0** | **0.8** | 3.2 | **5.0** | 15.4 | 3.0 | 18.4 | 6.1 |
| Spearman-HiE-PCA | 37.2 | 26.9 | 9.4 | 0.3 | **0.8** | 33.4 | **5.0** | 61.7 | 13.0 | 32.3 | 22.0 |
| Spearman-HiE-pPCA | 36.3 | 27.5 | 12.8 | 0.3 | 3.2 | 32.5 | **6.2** | 59.1 | 12.6 | 30.6 | 22.1 |

We observe that dimension reductions obtained using t stochastic neighbor embedding (tSNE) (van der Maaten and Hinton, 2008) perform better (on average) than those obtained with principal component analysis (PCA) or probabilistic PCA. Interestingly, we find that (dis)similarities in the reduced-dimensional space perform almost always better than in the original (dis)similarities (see Table 3). The main exception is RAFSIL2, which performs better using original similarities. We again see that approaches designed for scRNA-seq typically out perform more generic methods, and RAFSIL1 and RAFSIL2 have lower NNE compared with SIMLR. We note that Spearman correlation on highly-expressed genes, followed by tSNE, has good average performance comparable with RAFSIL1/2 and SIMLR.

We also visualize results from similarity learning and dimension reduction in Supplementary Figure S1. We find clear differences in the inferred similarities between methods for some datasets (especially for Leng and Usoskin, but also for Buettner), and this is reflected in the respective two-dimensional projections. Overall RAFSIL and SIMLR are able to more clearly separate cell populations compared to Euclidean distance and Spearman correlation. Also, we note that the good performance of RAFSIL2 (in terms of NNE, see Table 3) is clearly reflected, probably most pronounced for the Leng dataset. Overall, this shows that RAFSIL2 can improve the visualization (and therefore discovery) of group/population structure in scRNA-seq data.

***RAFSIL can discover unwanted variation in scRNA-seq data.*** In practice, dimension reduction is typically used for exploratory data analysis, for instance to find group structure in the data that might correspond to novel (sub)populations of cells. However, it can also be a valuable tool for data quality control, for instance when color coding additional information about cells (covariates) in a two-dimensional projection of the data. Figure 1 demonstrates this approach. The first row depicts tSNE plots for the Usoskin dataset, with RAFSIL2 projections in the first two panels and SIMLR projections in panels three and four. Color coding each cell with biological labels (four principal neuronal types) we see a clear separation with both approaches (panels one and three), but with substantial structure inside each neuronal cell type. Panels two and three reveal that this structure is likely a technical artifact. In these panels we color code the cells according to a technical variable (different cell picking sessions). For both approaches, RAFSIL and SIMLR, we clearly see that the perceived sub-structure in different neuronal types can largely be explained by the picking session. For clarity, we have annotated one cell type (tyrosine hydroxylase containing neurons) in panels one and three with the colors of the technical annotation in the adjacent plot that correlate with prominent sub-clusters.

The second row in Figure 1 is set up in the same way, just this time using the data of Kolodziejczyk *et al.* (2015). Here the biological color coding corresponds to different culturing conditions of mouse embryonic stem cells, while the technical variable denotes different sequencing chips. In the RAFSIL representation (panels one and two) we again see sub-structure in the biological annotation that perfectly corresponds to technical annotation (different sequencing chips). For this dataset SIMLR also recapitulates the biological group structure (panel three), but does not pick up the presence of confounding technical variation (panel four).

In summary, Figure 1 shows that RAFSIL can detect unwanted technical variation in scRNA-seq data, also in cases where other methods do not. We note that in both publications the authors corrected for batch effects, and we have used the uncorrected data for these analyses. In practice, this type of approach is mainly useful to assess if corrections for known technical factors are successful, or to rule out that discovered group structure corresponds to known covariates. Also, we note that the choice of dimension reduction technique plays a role in these analyses; for instance, when using PCA instead of tSNE things become considerably less clear (data not shown). However, this is not unexpected given the good performance of tSNE as a dimension reduction method (see Table 3).

#### 3.2.3 Clustering

***Random forest based similarities accurately recapitulate annotated cell populations.*** Next, we explored the performance of RAFSIL1/2 in terms of cell clustering, which is commonly used to discover population/group structure in scRNA-seq data and constitutes an essential step for most analyses in this field. To do so, we used the dissimilarities learned by RAFSIL1/2 in two ways: *(i)* to perform hierarchical clustering of cells (HC), and *(ii)* as input for k-means clustering (KM), taking each cell as a vector of its similarities with all cells in the dataset. We use the adjusted rand index (ARI) and normalized mutual information (NMI) as quality measures (see Section 2.4.3 for details), and results are summarized in Table 4. As before, we compared RAFSIL1/2 to SIMLR and Spearman correlation, and added the direct application of HC and KM to the expression data (Data– in Table 4). Because there are more methods for clustering scRNA-seq data than for similarity learning, we added additional comparisons with SC3, SINCERA and pcaReduce that do not implement similarity learning but perform clustering.

**Table 4:**
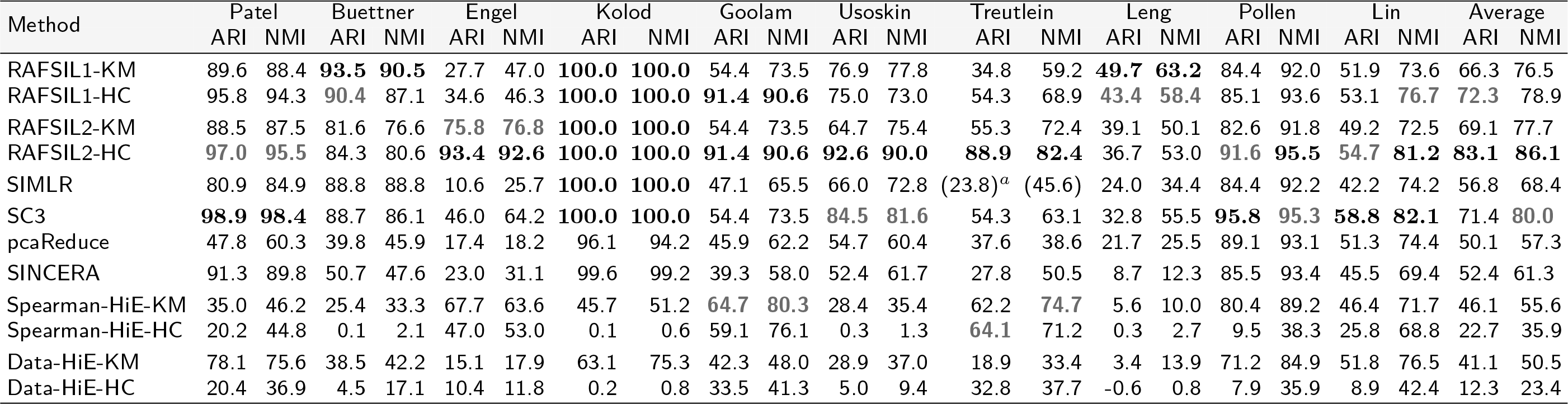
ARI and NMI values for clustering methods across ten data sets (in percent, higher is better). KM = k-means, HC = hierarchical clustering. “ Parentheses indicate that SIMLR was run with different parameters for this dataset.

We see that domain-specific methods for scRNA-seq clustering perform well, and that RAFSIL2 (using hierarchical clustering) has the best average performance, with SC3 and RAFSIL1-KM performing better for some datasets (Buettner, Patel and Leng). Interestingly, k-means clustering appears to perform better when directly applied to the data or in the context of Spearman correlation, while hierarchical clustering works better for random forest derived distances.

***Dimension reduction improves clustering.*** Motivated by our previous result of decreased NNE for reduced-dimension embeddings obtainable by tSNE, we applied clustering after dimension reduction for the methods we studied before (clustering-only approaches do not allow for dimension reduction). Results are summarized in Table 5, please see Section 2.4.3 for details on the Methods. Like before, we observe an overall better performance of clustering when using data with reduced dimensionality, again with the exception of RAFSIL2, which performs better in high dimensions. Also, comparing clustering results with similarity learning results, we find that using the original dissimilarity matrix RAFSIL2 had the smallest NNE and also the best clustering performance; for reduced dimensions, RAFSIL1 has the smallest NNE and also shows the best clustering performance. We finally note that the fact that RAFSIL2 performs worse that RAFSIL1 in this scenario is driven by its poor performance on the Kolod dataset. This relates to our previous discussion of Figure 1: batch effect removal may not have been successful for this dataset, and RAFSIL2’s clustering performance reflects the situation depicted in the first panel of the second row in, where cell groupings induced by cell picking session dominate biological variation.

**Table 5:** ARI and NMI values for clustering methods across ten data sets after dimension reduction (in percent, higher is better). ^α^ Parentheses indicate that SIMLR was run with different parameters for this dataset.

| Method | Patel |  | Buettner |  | Engel |  | Kolod |  | Goolam |  | Usoskin |  | Treutlein |  | Leng |  | Pollen |  | Lin |  | Average |  |
| --- | --- | --- | --- | --- | --- | --- | --- | --- | --- | --- | --- | --- | --- | --- | --- | --- | --- | --- | --- | --- | --- | --- |
|  | ARI | NMI | ARI | NMI | ARI | NMI | ARI | NMI | ARI | NMI | ARI | NMI | ARI | NMI | ARI | NMI | ARI | NMI | ARI | NMI | ARI | NMI |
| RAFSIL1-tSNE-KM | 96.8 | 95.7 | 93.5 | 90.5 | 44.7 | 55.9 | 100.0 | 100.0 | 54.4 | 73.5 | 61.7 | 71.5 | 33.8 | 58.2 | 46.4 | 61.1 | 89.0 | 94.9 | 48.6 | 74.1 | 66.9 | 77.5 |
| RAFSIL1-tSNE-HC | 93.4 | 91.8 | 93.5 | 90.5 | 26.6 | 46.3 | 100.0 | 100.0 | 54.4 | 73.5 | 64.6 | 75.8 | 54.8 | 70.9 | 46.6 | 62.4 | 89.2 | 94.9 | 50.1 | 76.3 | 67.7 | 78.2 |
| RAFSIL2-tSNE-KM | 97.5 | 96.3 | 87.3 | 83.1 | 26.6 | 46.3 | 34.9 | 41.7 | 54.4 | 73.5 | 65.5 | 77.1 | 55.0 | 72.4 | 48.7 | 60.0 | 88.0 | 93.3 | 42.1 | 71.2 | 60.0 | 71.5 |
| RAFSIL2-tSNE-HC | 97.5 | 96.3 | 87.5 | 85.0 | 24.8 | 45.1 | 30.9 | 38.9 | 54.4 | 73.5 | 65.9 | 78.5 | 55.8 | 72.5 | 30.9 | 46.2 | 87.5 | 93.3 | 48.8 | 73.6 | 58.4 | 70.3 |
| SIMLR-tSNE-KM | 90.8 | 89.6 | 88.8 | 88.8 | 10.6 | 25.7 | 100.0 | 100.0 | 47.1 | 65.5 | 66.0 | 73.4 | (27.3)<sup>a</sup> | (30.0) | 47.1 | 65.5 | 82.4 | 90.5 | 41.3 | 71.8 | 60.1 | 70.1 |
| SIMLR-tSNE-HC | 80.9 | 84.9 | 88.8 | 88.8 | 10.6 | 25.7 | 100.0 | 100.0 | 47.1 | 65.5 | 66.0 | 73.4 | (40.7) | (41.7) | 47.7 | 65.5 | 72.5 | 88.4 | 42.1 | 74.2 | 59.6 | 70.7 |
| Data-tSNE-KM | 71.5 | 72.2 | 33.4 | 33.0 | 18.0 | 18.9 | 92.6 | 90.0 | 35.8 | 52.6 | 84.9 | 80.2 | 31.6 | 55.1 | 16.7 | 26.4 | 82.3 | 88.8 | 54.4 | 78.5 | 52.1 | 59.6 |
| Data-tSNE-HC | 66.4 | 67.3 | 25.6 | 29.7 | 24.1 | 29.3 | 59.2 | 63.4 | 45.9 | 62.7 | 80.4 | 74.5 | 40.5 | 52.7 | 1.8 | 9.5 | 94.3 | 93.4 | 55.0 | 77.5 | 49.3 | 56.0 |
| Pearson-tSNE-KM | 88.5 | 86.2 | 29.2 | 33.6 | 28.9 | 35.4 | 100.0 | 100.0 | 58.2 | 74.0 | 64.9 | 66.6 | 40.2 | 62.0 | 6.9 | 10.9 | 78.6 | 91.1 | 48.1 | 73.0 | 54.4 | 63.3 |
| Pearson-tSNE-HC | 87.5 | 85.3 | 27.3 | 35.6 | 33.5 | 51.1 | 100.0 | 100.0 | 48.5 | 71.2 | 63.6 | 66.0 | 53.3 | 65.2 | 14.7 | 17.2 | 84.3 | 92.8 | 42.9 | 72.7 | 55.5 | 65.7 |

***RAFSIL approaches yield robust clustering solutions.*** To assess the robustness of clustering solutions, we randomly excluded 10% of cells from each dataset and re-run each clustering approach 20 times. Figure 2 summarizes the results. We see substantial variability in the ARI for most datasets and most methods across re-sampling runs; in terms of performance as measured by ARI averaged across datasets, RAFSIL2 (with hierarchical clustering) performs best with SC3 coming in second. This is consistent with our previous results obtained with the full data (see Table 4). Next, we looked at variability and calculated the inter quartile range (IQR) across res-sampling runs for each method analyzing each dataset, and then averaged across datasets (aIQR). SC3 exhibits the most stable clustering solutions (5% aIQR); RAFSIL2-HC is a bit worse with 7% aIQR, but a bit better than SIMLR, which has 8% aIQR. The method pcaReduce performs worst in terms of stability with an aIQR of 14%. Overall we find that RAFSIL produces relatively stable clustering solutions with good ARI.

**Figure 2:**
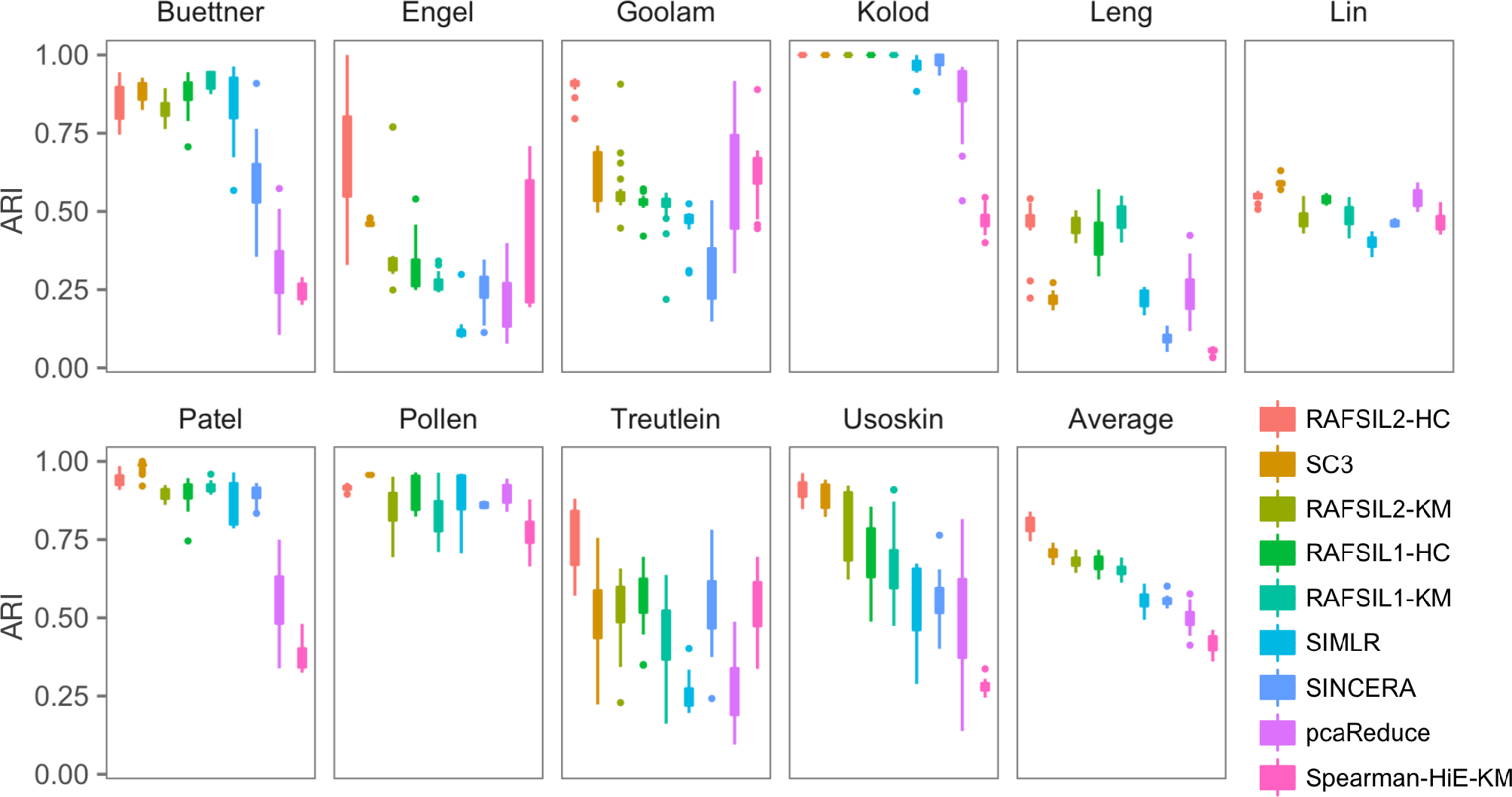
RAFSIL2 yields accurate and robust clustering solutions. Panels are box plots of the adjusted rand index (ARI) for ten datastes, across 20 instances of randomly sampling 90% of available cells. The panel labeled ‘Average’ represents the mean performance across all ten datasets. We see that RAFSIL2 followed by hierarchical clustering has the best performance, followed by SC3 and then the other RAFSIL-type methods. In terms of robustness SC3 performs best, while pcaReduce shows the highest variability. See Section 3.2.3 for a more detailed discussion. KM = k-means, HC = hierarchical clustering and HiE = highly expressed genes.

**Figure 3.**
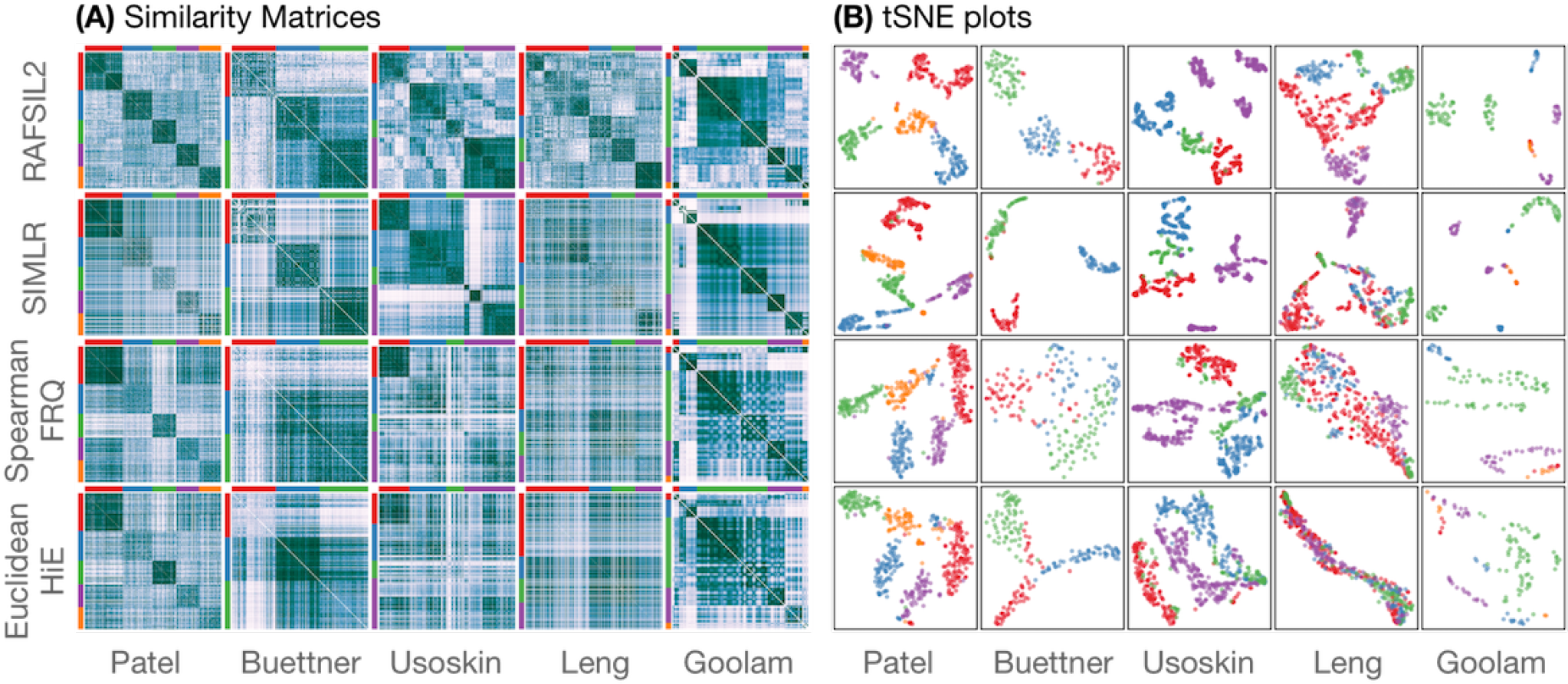
Figure S1. Similarity matrices and their visualization in two dimensions. Panel (A) shows similarity matrices obtained from four different methods (rows, see Section 2.4.3 in the main text) for four data sets (columns, see Table 1 in the main text)). Colors represent quantiles and darker colors stand for increasing similarity. Panel (B) shows corresponding tSNE plots, where each point represents a cell colored according to annotated cell types. The sub-panels in (B) capture differences in the similarity matrices on the left side, and we see that similarity learning affects visualization of SC-RNA-seq data sets. SIMLR and RAFSIL2 perform well on the Usoskin data set, and RAFSIL2 does especially well on the Leng dataset. Also see Section 3.2.2 in the main text.

***RAFSIL can estimate the number of populations in a scRNA-seq dataset.*** Here we ask whether RAFSIL can estimate the number of populations present in a scRNA-seq dataset. Briefly, we apply RAFSIL1/2 followed by hierarchical clustering (RAFSIL1/2-HC) and retrieve the corresponding series of cell partitions with increasing cluster numbers. To those we apply the Calinski-Harabasz criterion (Calinski and Harabasz, 1974), where each cell is described by its corresponding row in the scaled feature matrix ***F*** (see Section 2.2). We compared RAFSIL with SC3 and SINCERA in Table 6.

**Table 6:**
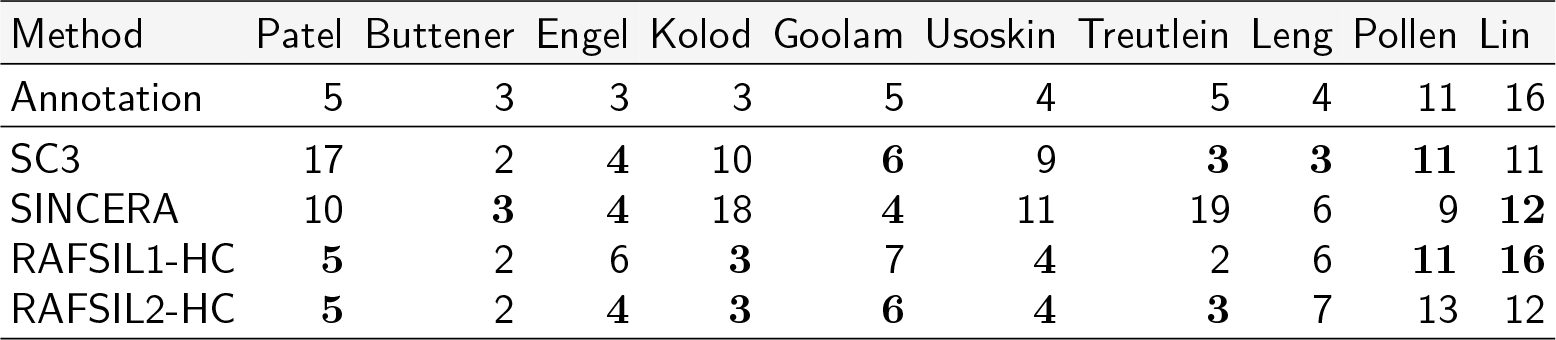
Number of clusters estimated by different methods for all data sets.

| Method | Patel | Buttner | Engel | Kolod | Goolam | Usoskin | Treutlein | Leng | Pollen | Lin |
| --- | --- | --- | --- | --- | --- | --- | --- | --- | --- | --- |
| Annotation | 5 | 3 | 3 | 3 | 5 | 4 | 5 | 4 | 11 | 16 |
| SC3 | 17 | 2 | 4 | 10 | 6 | 9 | 3 | 3 | 11 | 11 |
| SINCERA | 10 | 3 | 4 | 18 | 4 | 11 | 19 | 6 | 9 | 12 |
| RAFSIL1-HC | 5 | 2 | 6 | 3 | 7 | 4 | 2 | 6 | 11 | 16 |
| RAFSIL2-HC | 5 | 2 | 4 | 3 | 6 | 4 | 3 | 7 | 13 | 12 |

We find that RAFSIL1/2 perform well (RAFSIL2-HC is amongst the most accurate methods for the most datasets), but overall there is little difference between the approaches. ***Additional analyses.*** In addition to the analyses described above, we also compared our method to the neural network (NN) based approach of Lin *et al.* (2017). Lin et al. provide the data they used to assess their method, so we calculated performance metrics for RAFSIL1/2 and SC3 (without any gene filtering, to be consistent with the authors) and compared them to Table 2 from Lin *et al.* (2017). Results are shown in Table 7, where everything except the RAFSIL1/2 and SC3 lines has been taken from their publication. We see that the RAFSIL approaches (especially RAFSIL2) are competitive with the NN based approach, even though we do not make use of an supervised training phase.

**Table 7:**
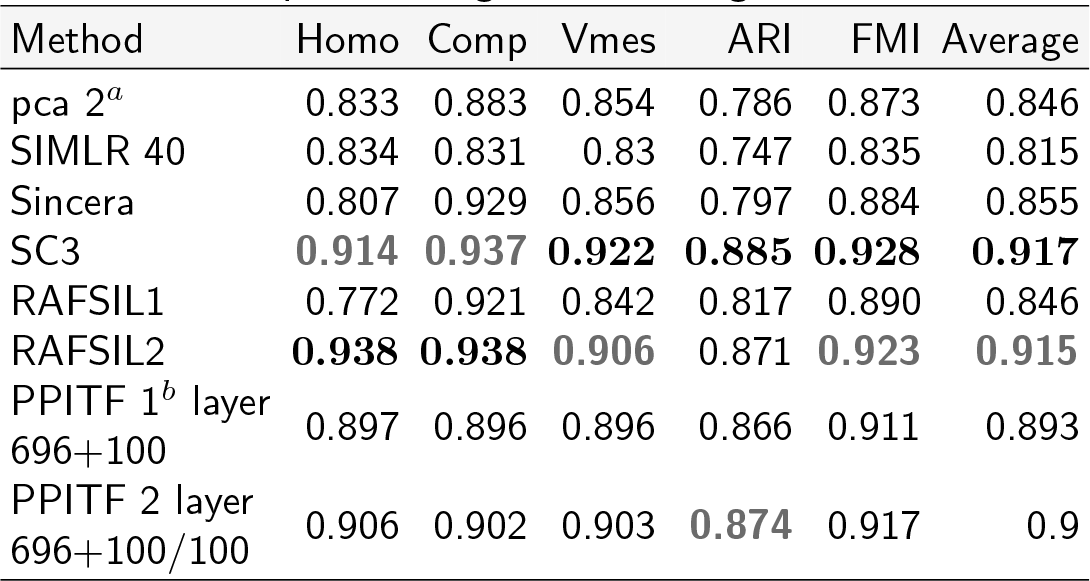
RAFSIL2 compared with a subset of the results reported by Lin *et al.* (2017). ARI: adjusted Rand index; Comp: completeness; FM: FowlkesMallows; Homo: homogeneity; Vmes: v-measure. *^a^*: pca with 2 principle components. *^b^*: PPITF uses prior biological knowledge to define a neural network architectures.

We also studied the clustering performance of RAFSIL1/2 performing only the feature construction step, and only the similarity learning step, respectively. Results are summarized in Table 8. We see that RAFSIL1/2 outperform these “reduced” approaches, highlighting the value of both of these steps in our approach. Nevertheless, feature construction alone followed by k-means clustering also performs well overall.

**Table 8:**
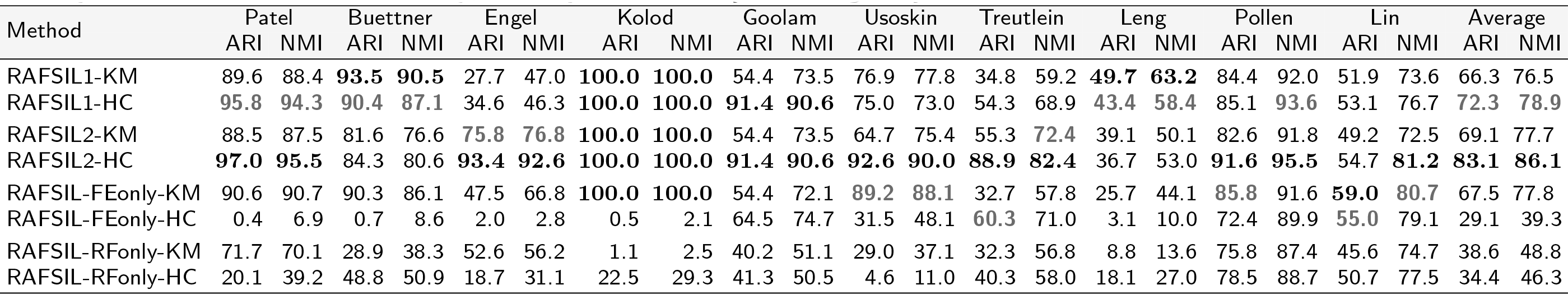
ARI and NMI values for clustering methods across ten data sets (in percent, higher is better). FEonly = feature construction only, RFonly = similarity learning only

## 4. Discussion and Conclusion

We have presented RAFSIL, a two-step approach for learning similarities between single cells based on whole transcriptome sequencing data. Accurately inferring such similarities is an important step in single cell RNA sequencing studies, because they form the basis for identification, visualization and interpretation of group structure. And reliable and accurate inference of group structure is necessary for discovery of new (sub)types of cells, for improved characterization and understanding of existing types of cells, for decoding the cellular composition healthy (and abnormal) tissue types, and more. We analyzed a diverse collection of datasets and show that RAFSIL performs well in similarity learning, on average outperforming SIMLR (to our knowledge the only other similarity learning approach geared specifically towards the scRNA-seq domain) as well as several generic approaches. In addition, the SIMLR algorithm requires a known (or pre-determined) number of clusters to calculate similarities, but reasonable estimates are not always available in practice. RAFSIL has no such requirement. We also show that RAFSIL similarities improve dimension reduction and data visualization, and that they can be used to discover unwanted technical variation in single cell RNA sequencing datasets. Finally, comparing clustering solutions obtained with RAFSIL similarities with state-of-the-art methods, we show that RAFSIL2 followed by hierarchical clustering is highly competitive, outperforming all other methods on average, and also individually on most datasets we studied.

Because RAFSIL implements a two-step procedure, first feature construction, and then similarity learning using random forests; it is flexible and easy to modify, expand and optimize. Our current feature construction step is a heuristic that reflects what we found to work well with scRNA-seq data we studied, but it is meant to be adapted as technology (and methodology) develops. For instance, including prior information about groups of genes (for example based on functional annotation databases) may improve performance. Likewise, we presented two strategies to apply random forests to unsupervised similarity learning (RAFSIL1 and RAFSIL2), but different approaches, perhaps more principled ones, can be imagined. Currently the time or of RAFSIL algorithms is comparable to methods like SC3 and SIMLR, and datasets with on the order of thousand cells can be analyzed without any problems. However, a truly large scale implementation for datasets with hundreds of thousands of cells (or more) would be desirable and is one of our future research directions.

Some limitations of our study include that while we compared RAFSIL extensively, our work is not exhaustive and results are restricted to the data we analyzed. However, we cover a variety of scRNA-seq technologies and computational approaches, and exhaustive comparisons considering all combinations of reasonable choices for gene filtering, dimension reduction, and clustering across many datasets quickly become infeasible. Along the same lines, we report that dimension reduction improves similarity learning and clustering, but only study projection into two-dimensional spaces (*k* = 2). While exploring larger choices for *k* might in principle be worthwhile for some methods, the fact that t stochastic neighbor embedding (tSNE) performed clearly best in our analysis might argue against it. The reason is that tSNE is known to perform well for projection into two to three dimensions, but runs into problems for higher ***k*** (van der Maaten and Hinton, 2008). Further on, we (and others) compare methods based on performance metrics like averages over adjusted Rand indexes (aARI) or average normalized mutual information. However, in our re-sampling experiment assessing robustness of clustering solutions (by repeatedly leaving out 10% of cells in a given dataset randomly) yields inter quartile ranges of the aARI between 5% and 14% (depending on the clustering method used). This implies that small performance differences are typically not robust to changing a small amount of cells in a dataset. While these values might be affected by the relatively small number of re-sampling runs (twenty), we believe it highlights the need for this type of analysis in the context of performance comparisons for single cell RNA-seq data methodology in general.

To summarize, we presented RAFSIL, a random forest based approach for similarity learning from single cell RNA sequencing data. We show that it performs well on a variety of datasets and believe it will be a useful tool for bioinformatics researchers working in this domain.

## Acknowledgements

We thank Dr. Abha S. Bais and Dr. Emmanuel Sapin for helpful discussions.

